# Open-source Tools for CryoET Particle Picking Machine Learning Competitions

**DOI:** 10.1101/2024.11.04.621608

**Authors:** Kyle I. Harrington, Zhuowen Zhao, Jonathan Schwartz, Saugat Kandel, Utz Ermel, Mohammadreza Paraan, Clinton Potter, Bridget Carragher

## Abstract

We are launching a machine learning (ML) competition focused on particle picking in cryo-electron tomography (cryoET) data, a crucial task in structural biology. To support this, we have created a comprehensive suite of open-source tools to develop resources for our competition, including copick for dataset management, napari plugins for interactive visualization, utilities for converting particle picks to segmentation masks, and PyTorch tools for custom dataset sampling. These resources streamline the processes of data handling, labeling, and visualization, allowing participants to focus on model development. By leveraging these tools, competitors will be better equipped to tackle the unique challenges of cryoET data and push forward advancements in particle picking techniques.

## 1 Introduction

We are hosting a machine learning competition to attract researchers focused on developing methods for identifying particle positions in 3D images acquired with cryo-electon tomography (cryoET). CryoET is a powerful technique for visual proteomics, enabling detailed exploration of biological systems at the molecular level. However, its application in large-scale experimentation is constrained by low throughput, particularly for identifying the 3D coordinates of proteins or macromolecular complexes within tomograms – crucial for achieving near-atomic resolutions with sub-tomogram averaging. This step of identifying particle positions is termed particle picking, the process of identifying and labeling individual particles within tomograms. Our competition is focused on supporting model development and evaluation for particle picking in cryoET data with an emphasis on identifying multiple particle types of varying sizes in experimental data.

Particle picking in cryoET is typically approached as an object detection or segmentation problem. Consequently, some of the most successful algorithms are based on model architectures such as YOLO [1], ResNet [2], and U-Net [3, 4]. Nevertheless, in practice, model performance can vary significantly depending on the dataset and the particle of interest.

To advance the field, several machine learning competitions have been organized with the aim of improving state-of-the-art techniques for particle picking. Unlike our competition, previous challenges

Preprint. Under review.in cryoEM have primarily focused on synthetic data. Past competitions in cryo-electron microscopy (cryoEM) and cryoET relied on synthetic data or focused on specific particles, such as ribosomes. For example, HREC 2020 for cryoEM [5], cryoET SHREC 2021 [6], and competitions on heterogeneity in cryoEM [7] have provided valuable benchmarks. Although these competitions have had a positive impact on the field, synthetic data is not ideal for training machine learning models. It often fails to capture the full range of distortions, artifacts, and noise often present in experimental data – which is critical for developing robust generalizable algorithms for real-world applications.

Our competition is designed to identify models that can: (1) maximize prediction performance when trained on a small number of annotated tomograms, and (2) perform robustly across a range of particle sizes. This setup reflects the typical scenario faced by cryoET researchers who need to minimize annotation effort while studying a diverse set of particles, often with only a few tomograms available for training. This paper outlines the open-source tools we have developed as resources to support participants in this and future cryoET machine learning competitions.

## 2 Data

To preserve the integrity of our competition, the data is thoroughly described in a manuscript that is currently under embargo until the official launch of the competition.

In the meantime, we have used a publicly available synthetic dataset with similar characteristics to demonstrate the open-source tools developed to support cryoET particle picking competitions. This dataset uses PolNet [8] to simulate 27 tomograms containing particles of varying sizes. This dataset is available as deposition CZCDP:10439 on the CZ cryoET Data Portal [9]. The dataset includes not only tomograms but also ground-truth picks in the copick format and binary segmentation masks for each particle species. An example tomogram is illustrated in Fig. 1 and is available here: https://cryoetdataportal.czscience.com/datasets/10439.

**Figure 1.**
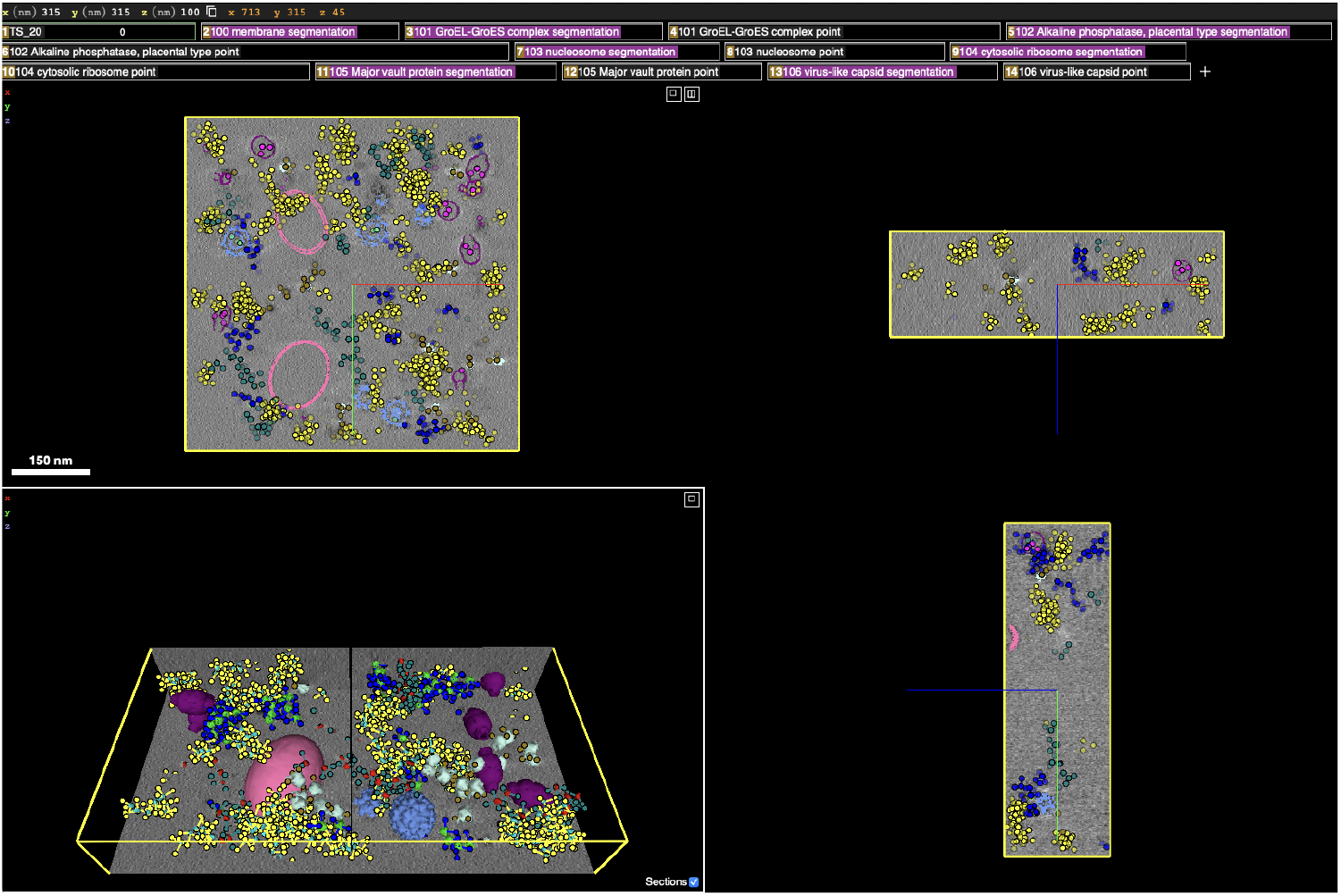
The orthogonal views of tomogram number 20 of the dataset 10439 through CryoET Data Portal.

**Figure 2.**
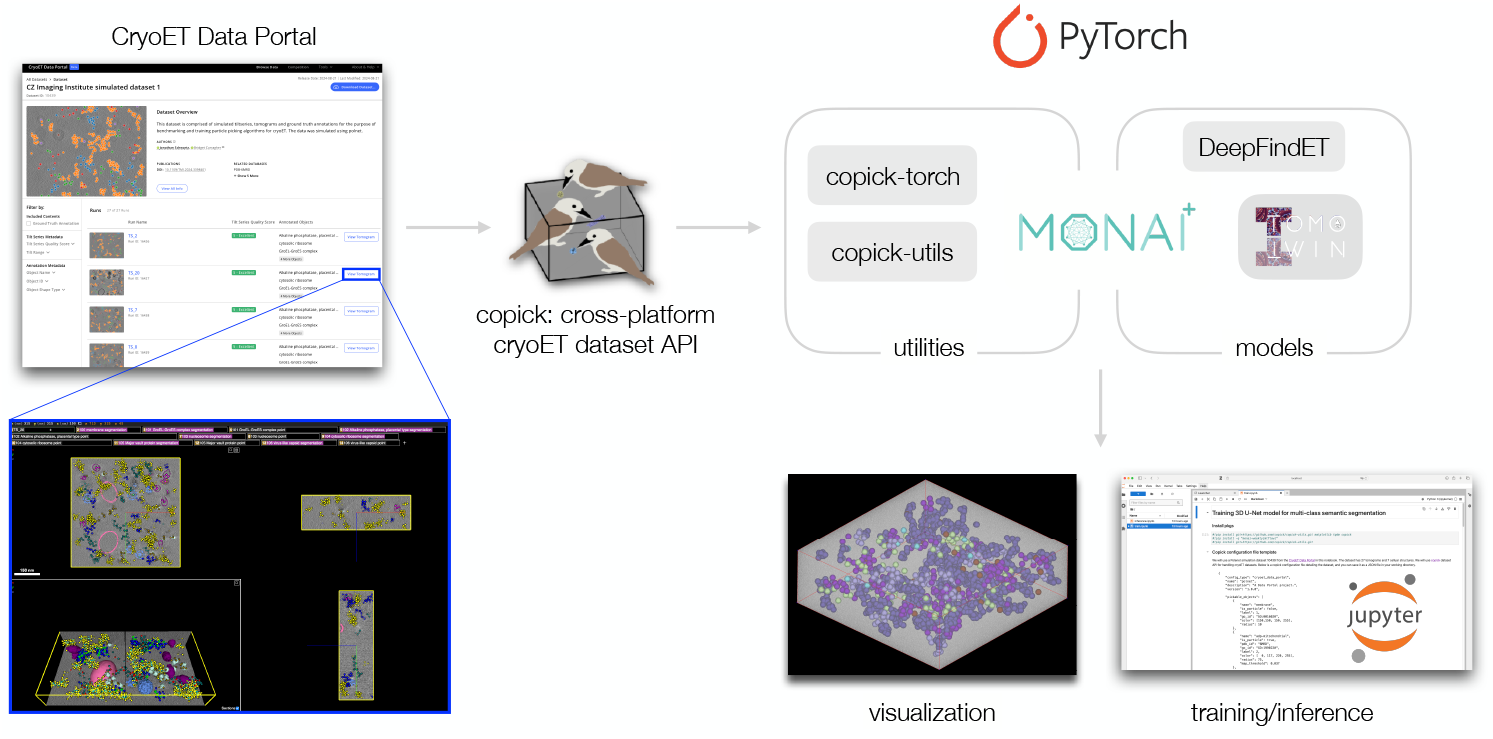
The software ecosystem that is used in resources for our cryoET ML Challenge.

## 3 Methods and Tools

In this paper, we present the suite of open-source tools we have used and created as resources to support participants in our cryoET particle picking competition. These tools cover a range of functionalities, including data management, visualization, data labeling, and PyTorch dataset loading and sampling.

For data management, we developed *copick* (https://github.com/copick/copick), a generic interface designed to handle cryoET datasets. To facilitate visualization, our example notebooks provide example code using *Matplotlib*, along with a custom *napari* plugin that enables users to visually browse and interact with *copick* projects. To streamline data labeling, we created *copick-utils* (https://github.com/copick/copick-utils), a collection of utility functions that includes conversion between point annotations and semantic segmentation masks. For PyTorch Dataset integration, we developed *copick-torch* (https://github.com/copick/copick-torch), which offers a PyTorch Dataset tailored for *copick* and sample code for visualizing sampling patterns within tomograms. Together, these tools allow us to create unified resources that simplify participation in cryoET particle picking competitions, helping researchers and developers focus on advancing particle picking techniques.

### Copick enables accessing, manipulating, and storing multimodal cryoET data—tomograms, collections of particle picks, segmentation masks, and feature sets

*Copick* provides a storageagnostic API that interfaces seamlessly with various data sources, including local, remote, and cloud-based platforms, such as the CZ CryoET Data Portal. The API is also used in other tools that are discussed in this paper, which makes it easier for our competition resources to enable easy reuse of competitors machine learning tools and pipelines across different data sources. We suggest that it is critical for competitors in cryoET particle picking competitions to be able to readily adapt and extend their datasets. An example *copick* project structure can be seen in the right-side panel of Fig. 3.

**Figure 3.**
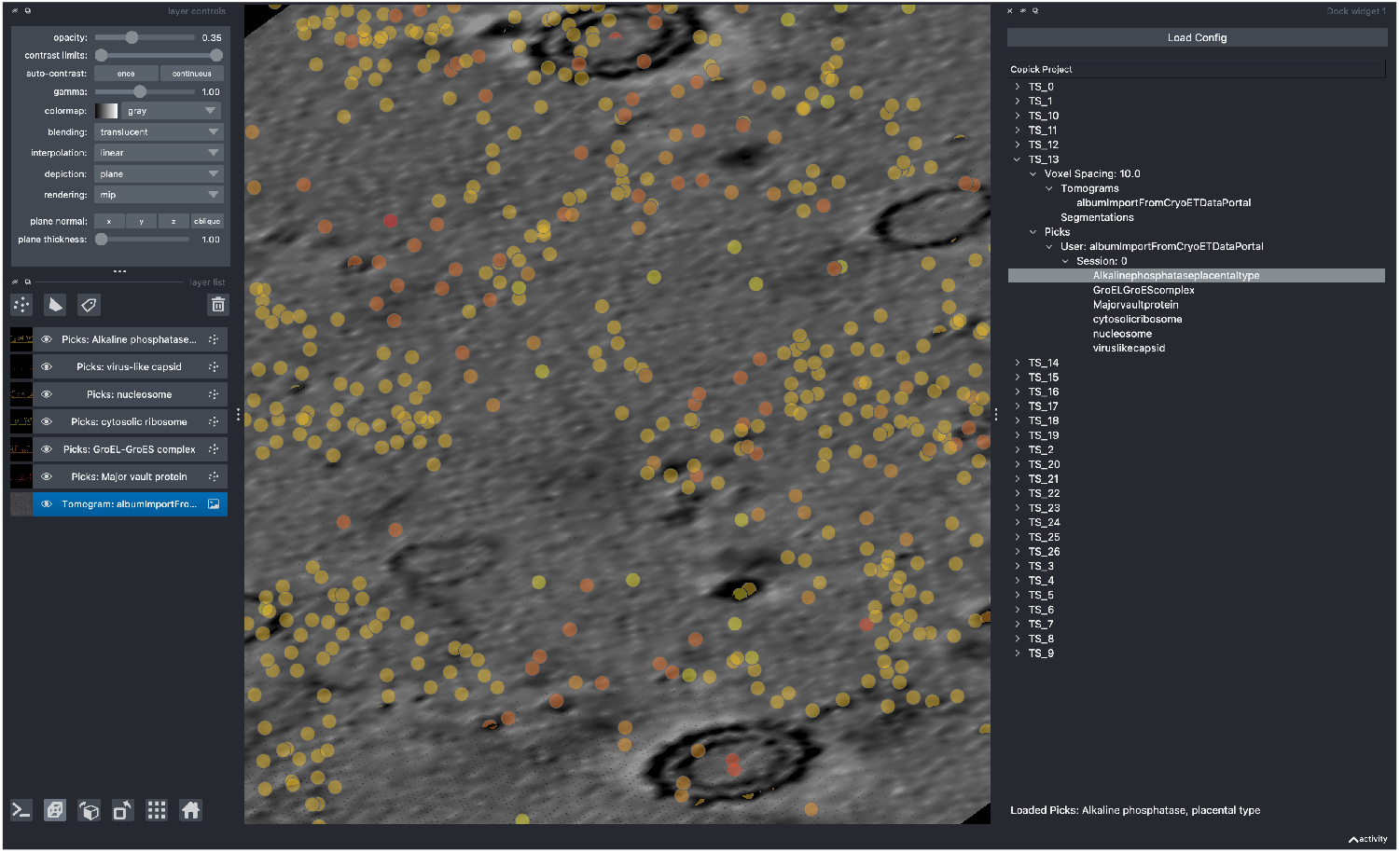
Example synthetic tomogram with picks overlaid in *napari-copick*.

### Visualizing data from a copick project can facilitate model debugging

A key step in model development and evaluation for image-based ML models is visualization. In our example notebooks we provide multiple methods for visualization of *copick* data: *Matplotlib* and *napari*. Furthermore, other more specialized tools like *ArtiaX* [10] can be leveraged for achieving a more holistic visualization from a biological perspective (e.g. visualization of particle structures in context).

To support interactive visualization we developed the *napari-copick* plugin which allows users to browse tomograms, particle picks, and segmentations directly from *copick* datasets. As a part of the visualization demos, we provide examples of visualizing PyTorch dataloaders, which are discussed in a later section. An example of visualizing a *copick* project using the *napari-copick* plugin is shown in Fig. 3.

### Converting between particle picks and segmentations

Models that approach particle picking as a supervised segmentation problem (e.g., *DeepFinder* [11] and *Topaz* [12]) require semantic segmentation masks for training, and generate segmentation masks as output. However, cryoET particle picking annotations are generally represented as single points for subtomogram averaging. To address this, segmentation based models often use particle coordinates to generate a segmentation mask, and then extract particle coordinates from segmentation masks. We provide routines for performing these operations on *copick* projects from the *copick-utils* Python module.

Our example notebooks provide our current guidance on creating these masks, but we encourage competitors to consider their own mask generation strategies. We create segmentation masks for particles by rasterizing, “painting,” each pick as a sphere that has a diameter based upon the particle type (for example, 50 %*±* 5 % the ribosome diameter). To convert segmentation masks back to particle picks we use connected components to convert semantic masks into instance masks, then perform watershed on each instance mask to split connected merged particles into separate instances, and finally take the centroid of each instance as a particle pick. An example figure showing particles and their corresponding rasterized masks is shown in Fig. 4.

**Figure 4.**
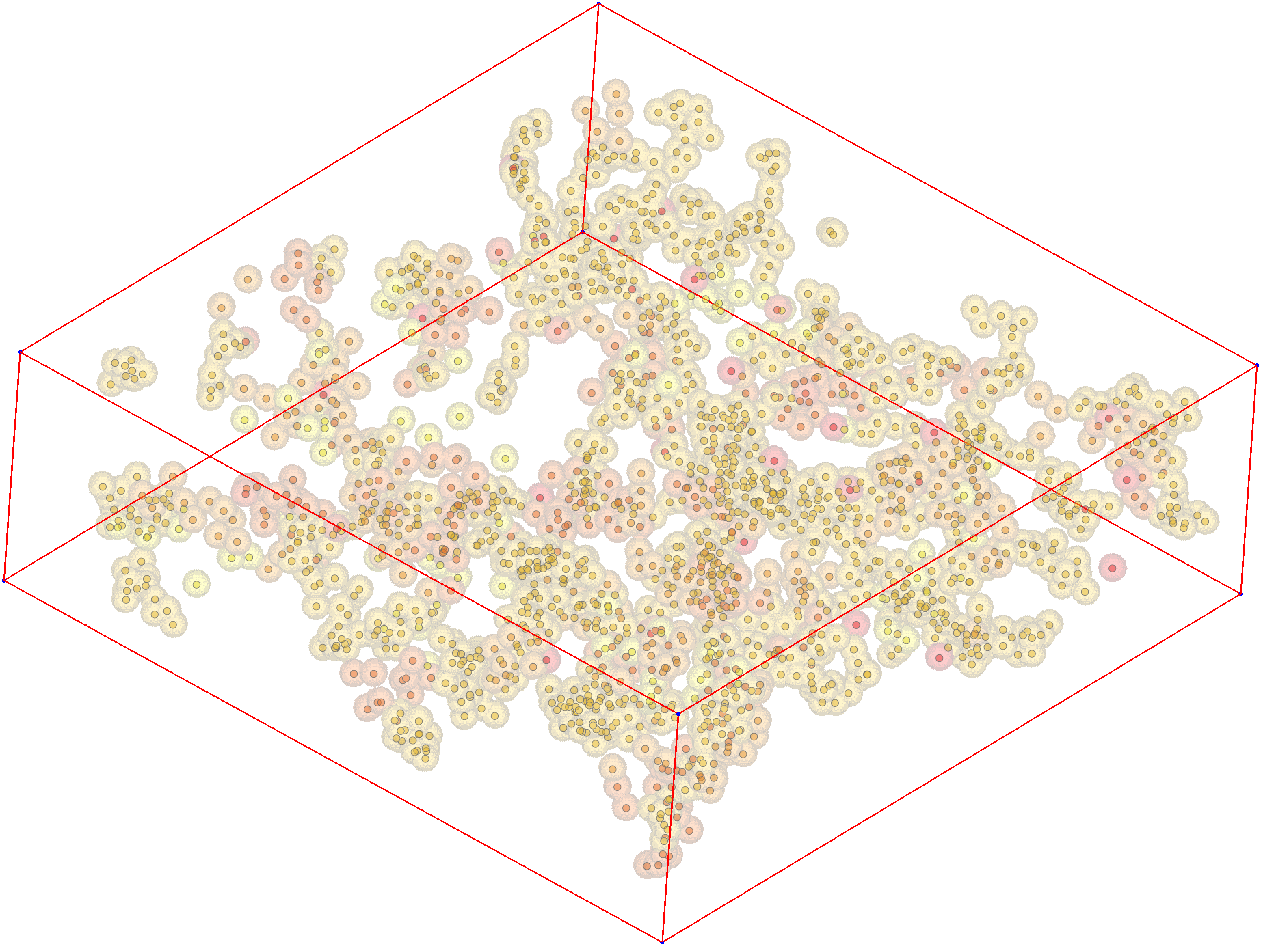
Picks from synthetic data and the output of segmentation from picks.

### Using and visualizing datasets with PyTorch

Innovations in data augmentation and dataset handling can have a significant impact on model performance. To facilitate investigations into *copick* datasets, we create example notebooks that use PyTorch-based datasets/dataloaders for *copick* from the *copick-torch* Python module, as well as tools to support the visualization of datasets (see Fig. 5). The *CopickDataset* for torch provides patchwise-sampling through the *morphospaces* python module, including the ability to tune segmentation density, striding, and other aspects of patch sampling. Figure 5b shows a visualization of a *CopickDataset* in *napari*.

**Figure 5.**
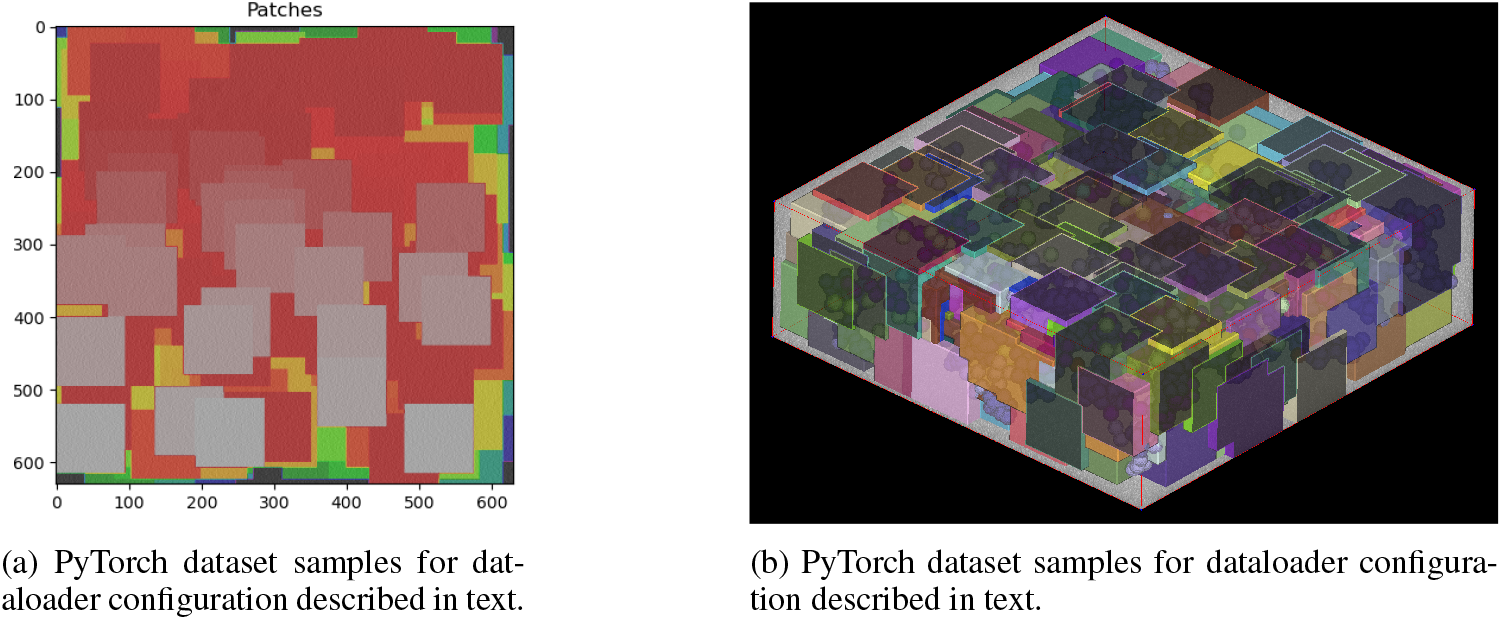
Two panels showing PyTorch dataset samples for different dataloader configurations.

### Example training and inference notebooks

We provide example notebooks utilizing the *copick* ecosystem for data handling (loading and preprocessing), allowing the challenge participants to focus on model development without having to worry about unfamiliarity with the cryoET data modality. Our core notebook is provided for using a 3D U-Net with PyTorch via MONAI [13]. This notebook and model is designed to be compatible with all the tools described in this paper. MONAI also provides additional models, such as ResUNet and Swin-UNETR, and our notebooks can be readily adapted to using these models as well. We provide example notebooks (the MONAI 3D U-Net, TomoTwin, and DeepFindET) in this repository https://github.com/czimaginginstitute/2024_czii_mlchallenge_notebooks.

### Getting more data for training

Our competition provides a limited amount of experimental tomograms for training data, but we supplement this challenge with by providing example synthetic data, tools to support generating synthetic data, and tools for accessing the CZ CryoET Data Portal. Generated synthetic data has the benefit of not only controlling the ground truth for particle locations, but the complete segmentation mask. We leverage PolNet [8] to generate tomograms and use a generic album-based [14] tool for tomogram generation https://copick.github.io/copick-catalog/polnet/generate-copick-project. *Copick* also interfaces with data portals, which can serve as another source of experimental data for supplementing model training.

## 4 Discussion and Conclusion

We have described a collection of open-source tools that we have leveraged to create example notebooks for cryoET particle picking competitions. These tools encompass a range of functionalities, including dataset management, automated data annotation generation, example model implementations, visualization, and synthetic data generation. The examples and notebooks presented in this paper demonstrate the end-to-end process of data exploration to model development and generation of predictions. By offering these resources, we aim to empower a broad audience, enabling participation in cryoET particle picking competitions with minimal prior experience in the domain.

## Acknowledgments and Disclosure of Funding

CZ Imaging Institute is made possible with support from the Chan Zuckerberg Initiative (CZII-2023–327779).

## References

[1] Thorsten Wagner, Felipe Merino, Markus Stabrin, Toshio Moriya, Claudia Antoni, Amir Apel-baum, Philine Hagel, Oleg Sitsel, Tobias Raisch, Daniel Prumbaum, et al. Sphire-cryolo is a fast and accurate fully automated particle picker for cryo-em. Communications biology, 2(1):218, 2019.

[2] Gavin Rice, Thorsten Wagner, Markus Stabrin, Oleg Sitsel, Daniel Prumbaum, and Stefan Raunser. Tomotwin: generalized 3d localization of macromolecules in cryo-electron tomograms with structural data mining. Nature methods, 20(6):871–880, 2023.

[3] Tristan Bepler, Kotaro Kelley, Alex J Noble, and Bonnie Berger. Topaz-denoise: general deep denoising models for cryoem and cryoet. Nature communications, 11(1):5208, 2020.

[4] Emmanuel Moebel, Antonio Martinez-Sanchez, Lorenz Lamm, Ricardo D Righetto, Wojciech Wietrzynski, Sahradha Albert, Damien Larivie’re, Eric Fourmentin, Stefan Pfeffer, Julio Ortiz, et al. Deep learning improves macromolecule identification in 3d cellular cryo-electron tomograms. Nature methods, 18(11):1386–1394, 2021.

[5] Ilja Gubins, Marten L Chaillet, Gijs van Der Schot, Remco C Veltkamp, Friedrich FoÖrster, Yu Hao, Xiaohua Wan, Xuefeng Cui, Fa Zhang, Emmanuel Moebel, et al. Shrec 2020: Classification in cryo-electron tomograms. Computers & Graphics, 91:279–289, 2020.

[6] Ilja Gubins, Marten L. Chaillet, Gijs van der Schot, M. Cristina Trueba, Remco C. Veltkamp, Friedrich FoÖrster, Xiao Wang, Daisuke Kihara, Emmanuel Moebel, Nguyen P. Nguyen, Tommi White, Filiz Bunyak, Giorgos Papoulias, Stavros Gerolymatos, Evangelia I. Zacharaki, Konstantinos Moustakas, Xiangrui Zeng, Sinuo Liu, Min Xu, Yaoyu Wang, Cheng Chen, Xuefeng Cui, and Fa Zhang. SHREC 2021: Classification in Cryo-electron Tomograms. In Silvia Biasotti, Roberto M. Dyke, Yukun Lai, Paul L. Rosin, and Remco C. Veltkamp, editors, Eurographics Workshop on 3D Object Retrieval. The Eurographics Association, 2021.

[7] Miro A. Astore, Geoffrey Wollard, David Silva-Sánchez, Wenda Zhao, Khanh Dao Duc, Nikolaus Grigorieff, and Pilar Cossio. The inaugural flatiron institute cryo-em heterogeneity communicty challenge, 2023.

[8] Antonio Martinez-Sanchez, Lorenz Lamm, Marion Jasnin, and Harold Phelippeau. Simulating the cellular context in synthetic datasets for cryo-electron tomography. bioRxiv, pages 2023–05, 2023.

[9] Utz Ermel, Anchi Cheng, Jun Xi Ni, Jessica Gadling, Manasa Venkatakrishnan, Kira Evans, Jeremy Asuncion, Andrew Sweet, Janeece Pourroy, Zun Shi Wang, et al. A data portal for providing standardized annotations for cryo-electron tomography. Nature Methods, pages 1–3, 2024.

[10] Utz H Ermel, Serena M Arghittu, and Achilleas S Frangakis. Artiax: an electron tomography toolbox for the interactive handling of s ub-tomograms in ucsf chimerax. Protein science, 31(12):e4472, 2022.

[11] Emmanuel Moebel, Antonio Martinez-Sanchez, Damien Larivie’re, Eric Fourmentin, Julio Ortiz, Wolfgang Baumeister, and Charles Kervrann. Deep learning improves macromolecules localization and identification in 3d cellular cryo-electron tomograms. bioRxiv, 2020.

[12] Tristan Bepler, Andrew Morin, Micah Rapp, Julia Brasch, Lawrence Shapiro, Alex J. Noble, and Bonnie Berger. Positive-unlabeled convolutional neural networks for particle picking in cryo-electron micrographs. Nature Methods, 2019.

[13] JM orge Cardoso, Wenqi Li, Richard Brown, Nic Ma, Eric Kerfoot, Yiheng Wang, Benjamin Murrey, Andriy Myronenko, Can Zhao, Dong Yang, et al. Monai: An open-source framework for deep learning in healthcare. arXiv preprint arXiv:2211.02701, 2022.

[14] Jan Philipp Albrecht, Deborah Schmidt, and Kyle Harrington. Album: a framework for scientific data processing with software solutions of heterogeneous tools. arXiv preprint arXiv:2110.00601, 2021.

